# Quantifying the effect of *A. thaliana* root on antibiotic production in the beneficial bacterium, *B. subtilis*

**DOI:** 10.1101/2021.06.03.446890

**Authors:** Harsh Maan, Omri Gilhar, Ziv Porat, Ilana Kolodkin-Gal

## Abstract

Beneficial and probiotic bacteria play an important role in conferring the immunity of their hosts against a wide range of bacterial, viral and fungal diseases. *B. subtilis* is a bacterium that protects the plant from various pathogens due to its capacity to produce an extensive repertoire of antibiotics. At the same time, the plant microbiome is a highly competitive niche, with multiple microbial species competing for space and resources, a competition that can be determined by the antagonistic potential of each microbiome member. Therefore, regulating antibiotic production in the rhizosphere is of great significance to eliminate pathogens and to establish beneficial host-associated communities.

In this work, we used *Bacillus subtilis* as a model to investigate the role of plant colonization in antibiotic production. Flow cytometry and Image-stream cytometry analysis supported the notion that *A. thaliana* specifically induced the transcription of the biosynthetic clusters for the non-ribosomal peptides surfactin, bacilysin and plipastatin and the polyketide bacillaene. This induction could be beneficial for the root as all clusters were shown to antagonize plant pathogens. Consistently, the root failed to induce PenP, a β-lactamase that increases only the fitness of the bacteria. Our results can be translated to improve the performance and competitiveness of beneficial members of the plant microbiome.

## Introduction

Rhizobacteria can promote plant growth directly by colonization of the root and exert beneficial effects on plant growth and development (Kloepper et al., 2004a). These bacteria are often designated plant growth-promoting rhizobacteria (PGPR). To date, various PGPR have been isolated, including various *Bacillus* species, *Burkholderia cepacia and Pseudomonas fluorescens*. These beneficial rhizobacteria can also confer fitness on their hosts by activating their immune system, and by antagonizing plant pathogens (Berg et al., 2017; Berg and Raaijmakers, 2018; Allaband et al., 2019). In addition to the direct promotion of plant growth, PGPR enhance the efficiency of fertilizers, and aid in degrading xenobiotic compounds (Adam et al., 2016; Berg et al., 2017).

Among growth promoting strains and biocontrol agens, *Bacillus subtilis* and its related species, such as *B. amyloliquefaciens, B. velezensis and B. mojavensis* are considered model organisms (Fan et al., 2017). In particular, the antimicrobial activity of *B. subtilis* was so far demonstrated against bacterial, viral and fungal soil-borne plant pathogens (Kloepper et al., 2004b; Nagorska et al., 2007). This activity is mediated largely by antibiotic production: approximately 5% of the *B. subtilis* genome is dedicated to the synthesis of antimicrobial molecules by non-ribosomal peptide synthetases (NRPS) or polyketide synthases (PKS/NRPS) (Stein, 2005; Ongena and Jacques, 2008; Kinsella et al., 2009). *In vitro* and *in planta* studies indicated the importance of four antibiotics for plant protection: surfactin, bacilysin, plipastatin and bacillaene (Hou and Kolodkin-Gal, 2020; Arnaouteli et al., 2021).

Surfactin, is a small cyclic lipopeptide induced during the development of genetic competence (Magnuson et al., 1994). The machinery for surfactin synthesis is encoded within the *srfAA– AB–AC–AD* operon (Kluge et al., 1988). Surfactin is a powerful surfactant with antibacterial (Gonzalez et al., 2011) and antifungal properties (Falardeau et al., 2013).

Bacilysin is a non-ribosomal dipeptide composed of L-alanine and amino acid L-anticapsin, that demonstrates antibacterial activity against a wide range of pathogens (Rajavel *et al*., 2009). Its synthesis is controlled mainly by the *bac* operon (*bacABCDE)* and is regulated by other enzymes such as thymidylate synthase, homocysteine methyl transferase and the oligopeptide permeases (Inaoka et al., 2003).

Fengycin/plipastatin (Tsuge et al., 2007) is a highly effective lipopeptide comprising 10 amino synthetized by five fengycin synthetases (*ppsA, ppsB, ppsC, ppsD, and ppsE*). Bacillaene and dihydrobacillaene (Butcher et al., 2007; Straight et al., 2007) are polyketides synthesized by an enzymatic complex encoded in the *pks* gene cluster (Butcher et al., 2007; Straight et al., 2007). This linear antimicrobial macrolides are synthesized by the PKSJLMNR cluster mega-complex (Straight et al., 2007).

Interestingly, we recently found that the plant host can enhance the efficiency of the killing of *Serratia plymuthica by B. subtilis* by inducing the synthesis of the antibiotic baceilaene (Ogran et al., 2019). These preliminary results raise the question on whether additional antibiotics that contribute to rhizocompatibility are induced by the plant to promote the colonization of preferred symbionts. To address this question, we considered the overall effect of the plant host in regulating four major antibiotics: surfactin, bacillaene, bacilysin and plipstatin, and an unrelated antibiotic resistance gene, the beta-lactamase PenP.

Our results indicate that the attachment with the root can specifically enhance antibiotic production (but not the β-lactamase PenP) and therefore affect the competitiveness of root-associated bacteria compared with their free-living counterparts.

## Results

Using flow cytometry, we examined the expression of transcriptional reporters P_*srfAA*_-yfp, P_*pksC*_-mKate, P_*bacA*_-gfp and P_*ppsA*_-gfp in static cultures. Specifically, we asked whether association with the plant would impact their expression. For this purpose, we compared bacteria cultured in liquid medium with bacteria attached to plant roots. The number of cells expressing each NRP and their respective mean intensity of fluorescence both increased for all these antibiotics’ promoters. In addition, the mean intensity of cells expressing from the *pps* and *pks* operons also increased (Figure 1).

**Figure 1:**
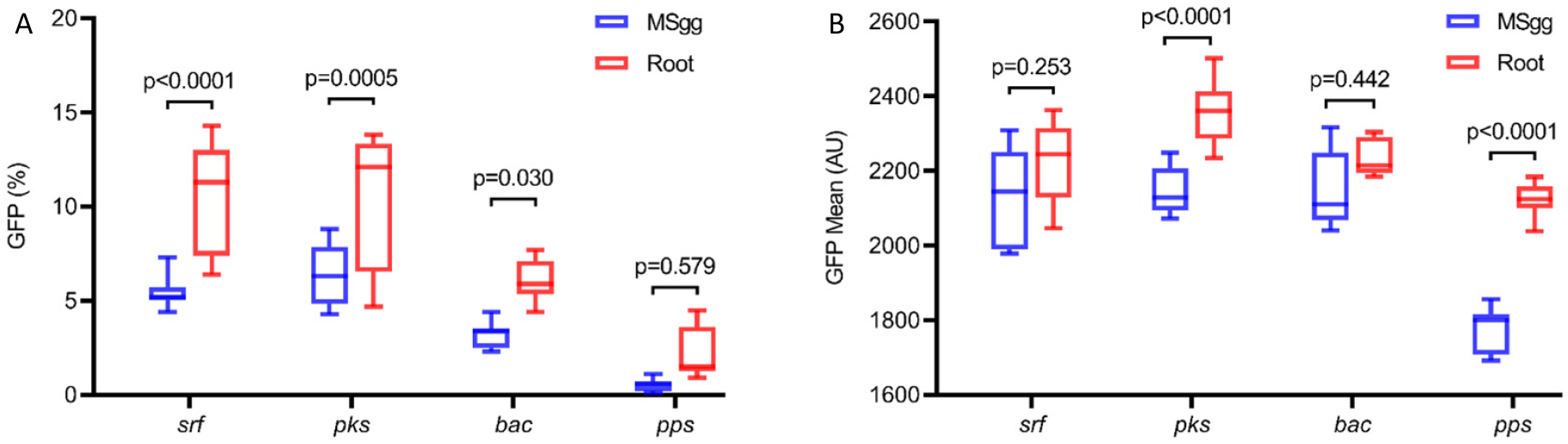
Indicated reporter strains for P_*srfAA*-_gfp (surfactin), P_*pksC*-_gfp (bacillaene), P_*bacA*-_gfp (bacilysin) and P_*ppsA*-_gfp (plipastatin), were analysed by flow cytometry for (A) positively expressing fluorescent populations or (B) the mean intensity of the fluorescent populations. Reporter stains were either grown in MSgg medium or in MSgg medium in presence of *A. thaliana* seedlings. Data were collected from 24 h post inoculation, 100,000 cells were counted. Graphs represent results from three independent experiments with n = 3/experiment (total n = 9/group). Statistical analysis was performed using Two-way ANOVA followed by Tukey’s multiple comparison post hoc testing. *P* < 0.05 was considered statistically significant.

Conventional flow cytometry allows accurate high-throughput quantification of fluorescence intensities, however it is less accurate for bacterial analysis. Higher fluorescence levels may be interpreted as higher expression levels, but also can result from larger bacterial size or small aggregates. Therefore, to increase our resolution into the manner by which antibiotic promoters respond to the attachment of the root we used Imaging Flow Cytometer. By collecting large numbers of digital images per sample and providing a numerical representation of image-based features, the ImageStreamX Mk II combines the per cell information content provided by standard microscopy with the statistical significance afforded by large sample sizes common to standard flow cytometry.

This allowed us to exclude most of the bacterial doublets or small aggregates, and calculate more accurately both the % of positive GFP cells, as well as bacterial cell length and GFP intensity normalized for cell size. This detailed analysis of antibiotic reporters demonstrated that in addition to the % of positive GFP cells for surfactin promoter, the distribution of GFP intensity in the population also changes (Figure 2). These effects were also observed for the transcription from the *bac*, (Figure 3), *pps (*Figure 4) and *pks* promoter (Figure 5). The impact of the plant on cell length was not robust, as it was not detected in all image-stream experiments (Figures 2-5). However, there were experimental sets where the plant significantly affected cell shape and reduced the length of *B. subtilis* cells (Figures 4 and 5).

**Figure 2:**
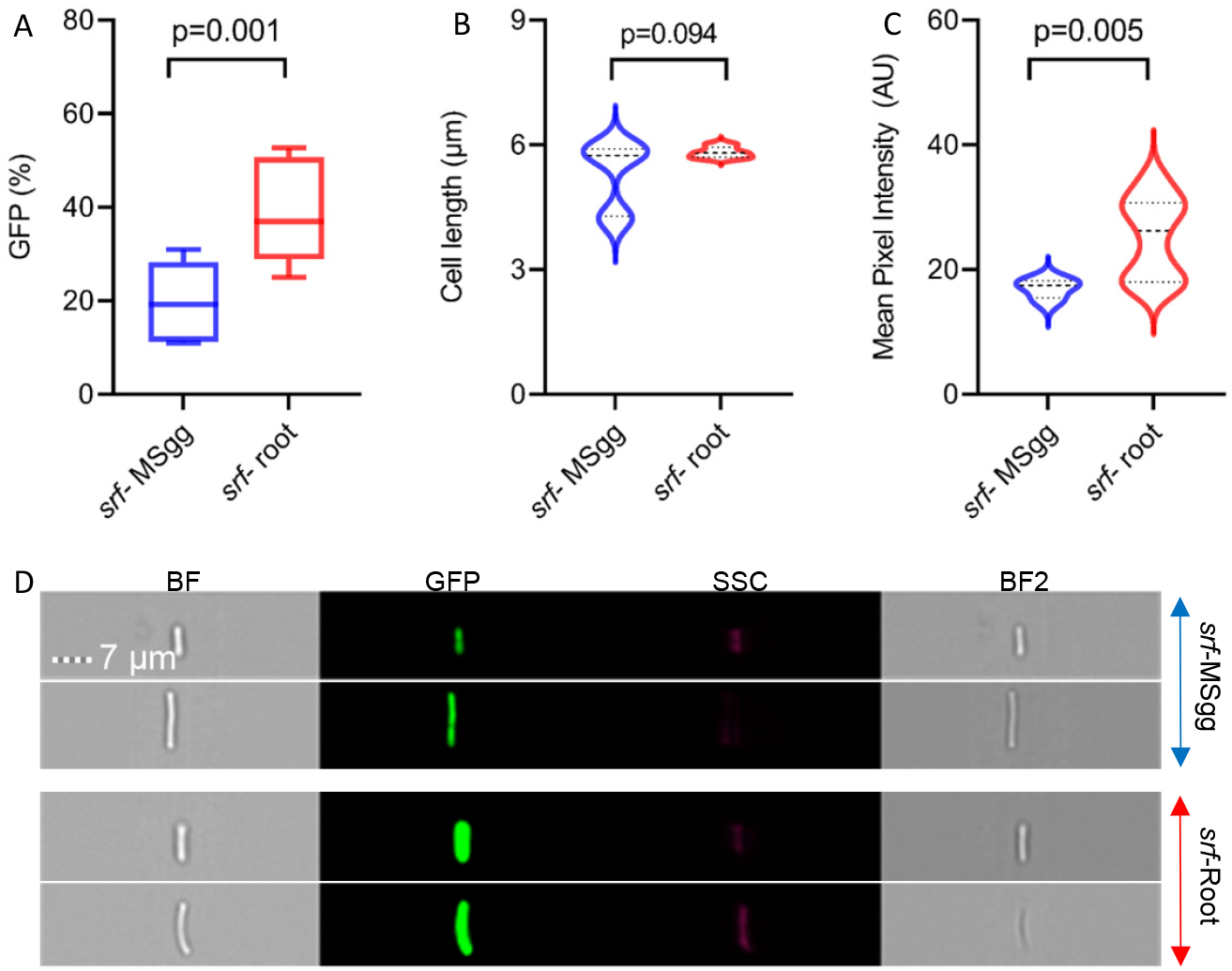
The expression of P_*srfAA*-_gfp (surfactin) attached to plant roots versus cultured in liquid MSgg. Bacteria expressing GFP under the control of the *srf* promoter were cultured in the absence or presence of *A. thaliana* seedlings. After 24h, bacteria colonizing the roots and bacteria cultured in liquid MSgg alone were collected. The percent of GFP expressing cells (**A**), cell length (**B**) and mean fluorescence intensity of the cells (**C**) were measured by Imaging Flow Cytometry and analyzed with IDEAS 6.3. (**D**) Representative bright field and fluorescent images related to expression of GFP in MSgg and on *A. thaliana* roots. Scale bar = 7 µm. Data were collected from 24 h post inoculation, 100,000 cells were counted. Graphs represent results from three independent experiments with n = 3/experiment (total n = 9/group). Statistical analysis was performed using unpaired t-test with Welch correction. *P* < 0.05 was considered statistically significant.

**Figure 3:**
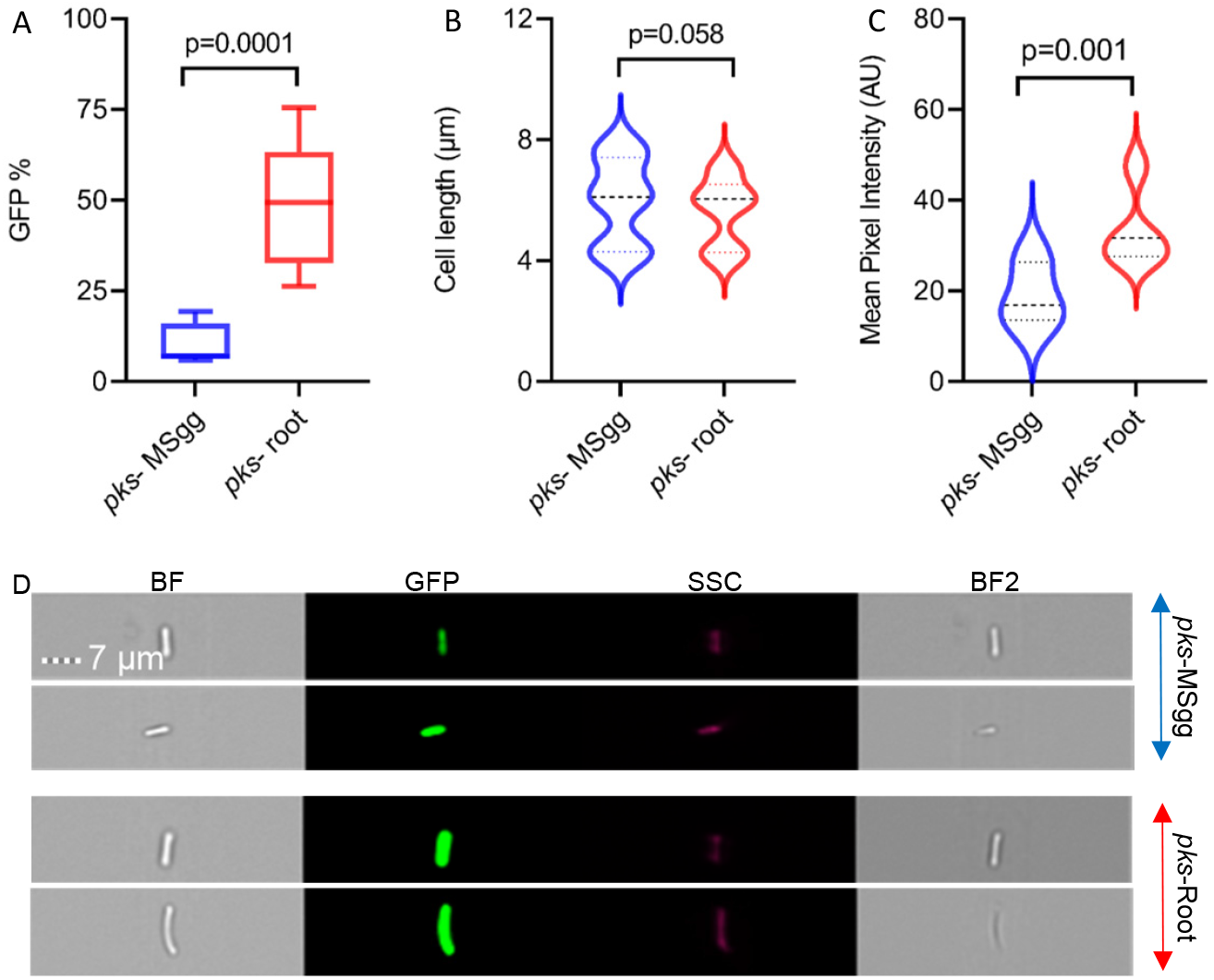
The expression of P_*pksC*-_gfp (bacillaene) attached to plant roots versus cultured in liquid MSgg. Bacteria expressing GFP under the control of the *pks* promoter were cultured in the absence or presence of *A. thaliana* seedlings. After 24h, bacteria colonizing the roots and bacteria cultured in liquid MSgg alone were collected. The percent of GFP expressing cells (**A**), cell length (**B**) and mean fluorescence intensity of the cells (**C**) were measured by Imaging Flow Cytometry and analyzed with IDEAS 6.3. (**D**) Representative brightfield and fluorescent images related to expression of GFP in MSgg and on *A. thaliana* roots. Scale bar = 7 µm. Data were collected from 24 h post inoculation, 100,000 cells were counted. Graphs represent results from three independent experiments with n = 3/experiment (total n = 9/group). Statistical analysis was performed using unpaired t-test with Welch correction. *P* < 0.05 was considered statistically significant.

**Figure 4:**
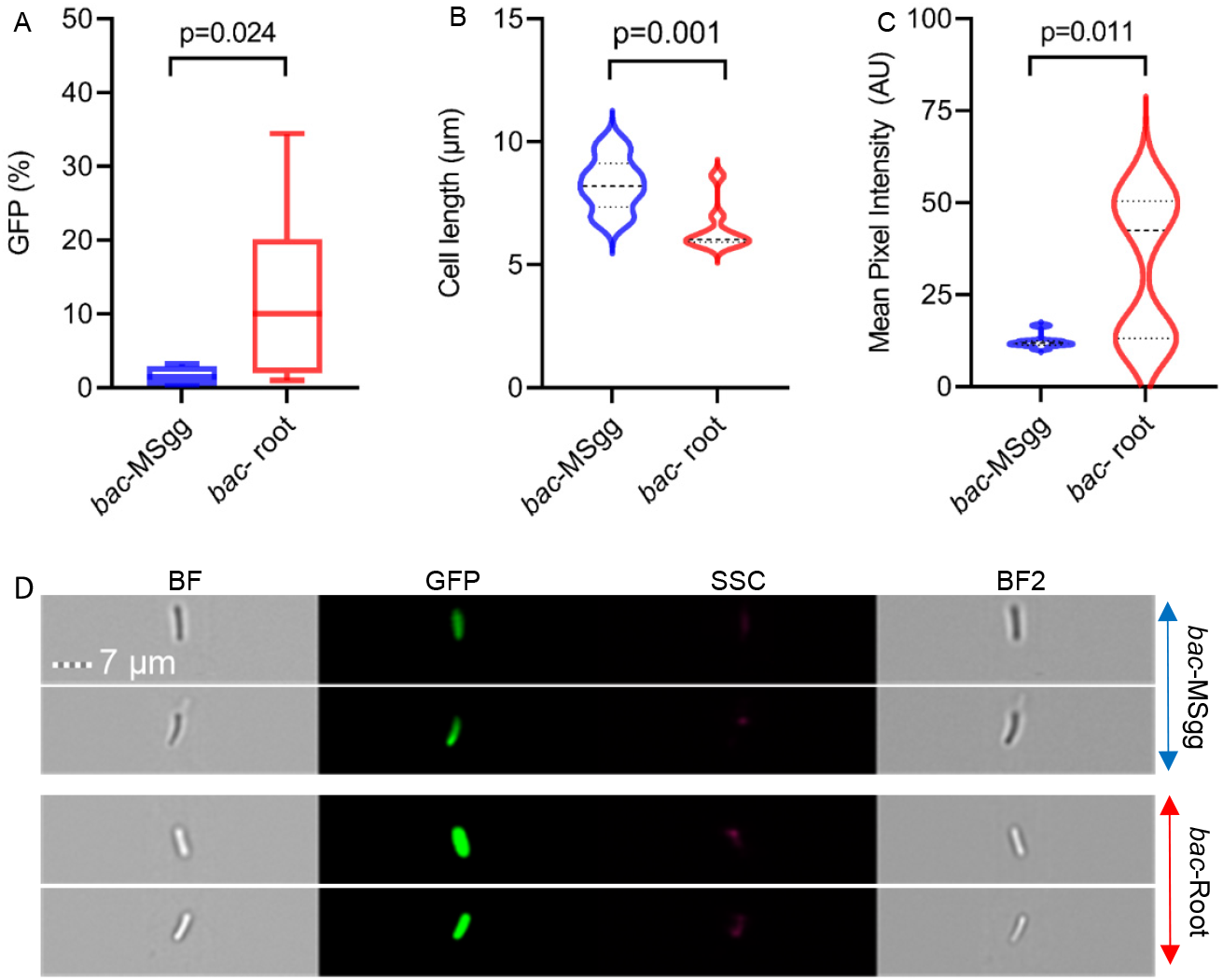
The expression of P_*bacA*-_gfp (bacilysin) attached to plant roots versus cultured in liquid MSgg. Bacteria expressing GFP under the control of the *bac* promoter were cultured in the absence or presence of *A. thaliana* seedlings. After 24h, bacteria colonizing the roots and bacteria cultured in liquid MSgg alone were collected. The percent of GFP expressing cells (**A**), cell length (**B**) and mean fluorescence intensity of the cells (**C**) were measured by Imaging Flow Cytometry and analyzed with IDEAS 6.3 (**D**) Representative brightfield and fluorescent images related to expression of GFP in MSgg and on *A. thaliana* roots. Scale bar = 7 µm. Data were collected from 24 h post inoculation, 100,000 cells were counted. Graphs represent results from three independent experiments with n = 3/experiment (total n = 9/group). Statistical analysis was performed using unpaired t-test with Welch correction. *P* < 0.05 was considered statistically significant.

**Figure 5:**
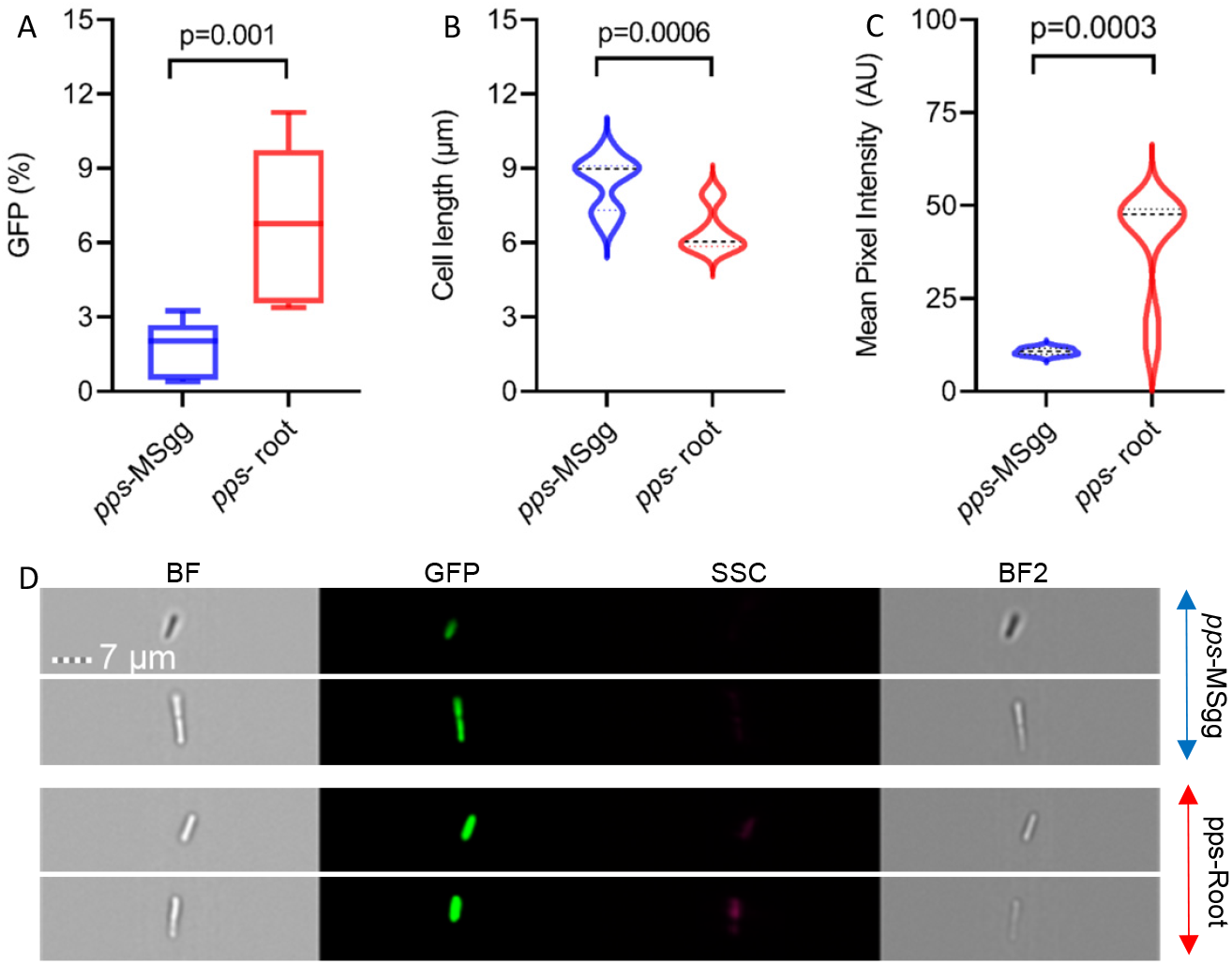
The expression of P_*ppsA*-_gfp (plipastatin) in bacteria attached to plant roots versus cultured in liquid MSgg. Bacteria expressing GFP under the control of the *pps* promoter were cultured in the absence or presence of *A. thaliana* seedlings. After 24h, bacteria colonizing the roots and bacteria cultured in liquid MSgg alone were collected. The percent of GFP expressing cells (**A**), cell length (**B**) and mean fluorescence intensity of the cells (**C**) were measured by Imaging Flow Cytometry and analyzed with IDEAS 6.3. (**D**) Representative brightfield and fluorescent images related to expression of GFP in MSgg and on *A. thaliana* roots. Scale bar = 7 µm. Data were collected from 24 h post inoculation, 100,000 cells were counted. Graphs represent results from three independent experiments with n = 3/experiment (total n = 9/group). Statistical analysis was performed using unpaired t-test with Welch correction. *P* < 0.05 was considered statistically significant.

To confirm that the impact of the plant root on antibiotic production could be due to a secreted we monitored the expression from P_*srfAA*_ and P_*pksC*_ fused to a luciferase reporter in the presence and absence of root secretions. The use of the unstable luciferase (an enzyme which degrades rapidly and therefore not accumulated (McLoon et al., 2011a) as a reporter allows us to monitor gene expression in real time by monitoring light production in a plate reader with an illuminometer. When grown on liquid defined medium, wild-type cells expressed luciferase from *srfAA* and *pksC* promoters robustly. However, while root secretions did not alter the kinetic of the expression, they were sufficient to significantly increase the intensity of the luciferase emission (Figure 6). These results suggest that the plant may secret metabolite that regulate the production of sufractin and bacillaene. Consistently, using confocal scanning laser microscopy, we could clearly confirm the expression of the *pks* promoter and on some extent, surfactin on cells attached with *A. thaliana* roots (Figure 7).

**Figure 6:**
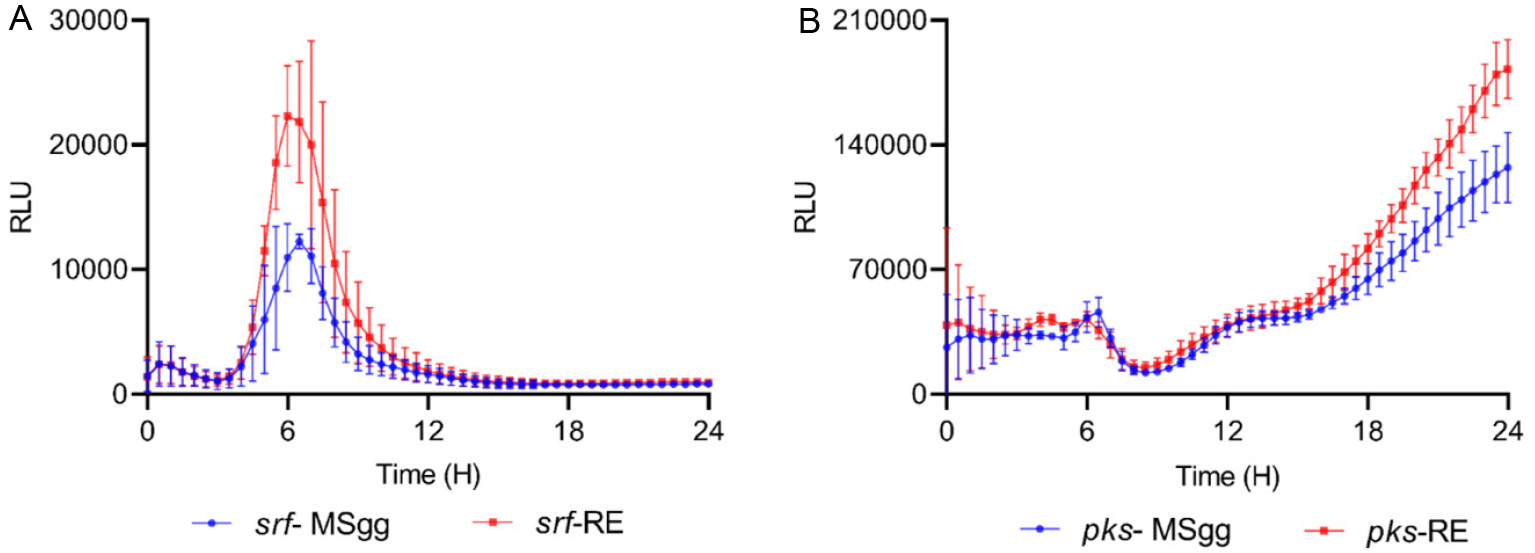
*A. thaliana* secretions increase the expression of (A) P_*srfAA*-_lux (surfactin) and (B) P_*pksC*-_ lux (bacilllane) in *B. subtilis* cells. Bacteria expressing luciferase under the control of the *srf* and *pks* promoters were cultured in *A. thaliana* secretions or in liquid MSgg and grown in a microplate reader for 24h. Graphs represent results from three independent experiments. Error bars represent ± SEM.

**Figure 7:**
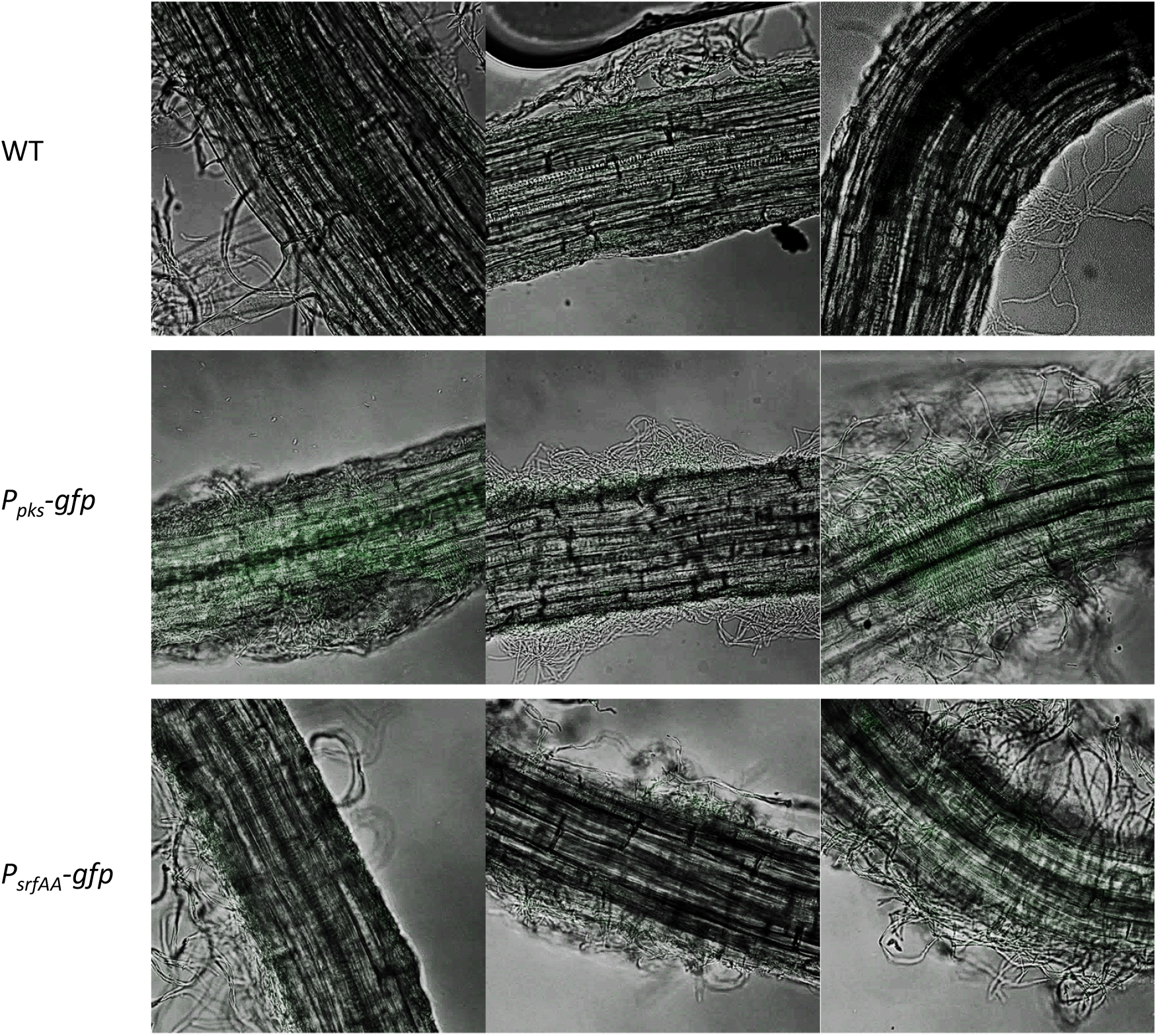
The expression of P_*pksC*-_gfp (bacillaene) and P_*srfAA*-_gfp (surfactin) on *A. thaliana* roots. Bacteria expressing GFP under the control of the *pks* and *srf* promoters were cultured in the presence of *A. thaliana* seedlings in MSgg medium. After 24h, the bacteria colonizing the roots were photographed with confocal microscope.

β-lactamases are enzymes which account for an additional layer of defence as they hydrolyse the β-lactam ring of β-lactams, thus inactivating the antibiotic before it reaches its target, the PBPs (Therrien and Levesque, 2000). Consistent with a hypothesis that the plant and tis secretions specifically regulate antibiotic production, the expression of the unrelated PenP promoter was not induced but rather slightly decreased in the presence of the plant (Figure 8).

**Figure 8:**
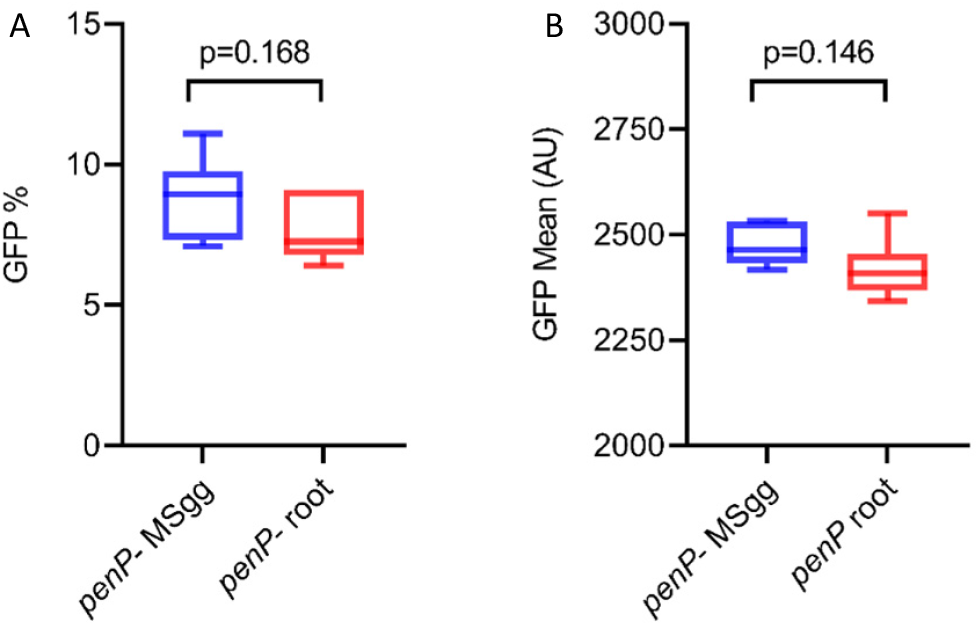
Indicated reporter strains for P_*penP*-_gfp was analysed by flow cytometry for (A) positively expressing fluorescent populations or (B) the mean intensity of the fluorescent populations. Reporter stains were either grown in MSgg medium or in MSgg medium in presence of *A. thaliana* seedlings.

## Discussion

Since *B. subtilis* was first described by Ferdinand Cohn in the late 1800s, it was shown to specialize in the production of metabolites (McLoon et al., 2011b). Many of the biosynthetic pathways for these metabolites are conserved either across the entire Bacillus genus or within specific phylogenetic clades. Fengycin, bacillaene, bacilysin and surfactin were essentially observed within the *B. subtilis* group (Hou and Kolodkin-Gal, 2020). This suggests that the different environmental niches inhabited by members of the *B. subtilis* clade may select for conservation of metabolites with distinct (or potentially redundant) beneficial functions.

Here, we found that the production of non-ribosomal peptides and polyketides was specifically activated during symbiotic interaction with *A. thaliana*. Our results demonstrated that both the root and its secretions increased the expression of four different biosynthetic clusters encoding for antibiotics with significance for *B. subtilis* competitiveness. In contrast, the PenP β-lactamase which deactivate β-lactam antibiotic was not induced by the host. This result is intriguing as the production of this enzyme promotes increased resistance of the bacterium in the competitive soil environment, containing β-lactam producers (Hou and Kolodkin-Gal, 2020), but does not contribute directly to the fitness of the plant. In contrast, each of the induced antibiotics were shown to grant protection to the plant versus fungal and bacterial pathogens.

Interestingly, we previously demonstrated that the interaction with the plant increases the capacity of *B. subtilis* to compete with *Serattia Plymuthica*, and our current results further indicate that the root is active regulator of the competitive interactions occurring on its roots (Ogran et al., 2019). The complexity of these antibiotic-host interactions suggests that *B. subtilis* biofilms can be considered a part of the plant microbiome, with the host actively promoting the establishment of the most beneficial bacterial community. Our findings provide a simple example of high-order interactions that shape microbiomes; the host modulates antibiotic production in the desired bacterial colonizers, providing the colonizers a clear advantage over less beneficial potential residents.

## Materials and Methods

### Strains and media

All strains used in this study are in Table S1. The strains were grown in LB broth (Difco), or MSgg medium(Branda et al., 2001) (5 mM potassium phosphate, 100 mM MOPS (pH 7), 2 mM MgCl2, 50 µM MnCl2, 50 µM FeCl3, 700 µM CaCl2, 1 µM ZnCl2, 2 µM thiamine, 0.5% glycerol, 0.5% glutamate, 50 µg/mL threonine, tryptophan and phenylalanine) (Branda et al., 2001). Solid LB medium contained 1.5% bacto agar (Difco).

### Plant Growth Conditions

Seeds of *A. thaliana* Col-0 were surface-sterilized and seeded on petri dishes containing Murashige and Skoog medium (4.4 g/L), PH 5.7, supplied with 0.5% (w/v) plant agar (Duchefa), 0.5% Sucrose (SigmaAldrich) and then stratified at 4°C for 2 days. The seeds were further transferred to a growth chamber (MRC) at 23°C in 12 h light/12 h dark regime.

### Extraction of plant secretions

Plant secretions were retrieved from 14-day-old *A. thaliana* seedlings cultured in 6 ml liquid MSgg of each well of a 6-well microplate (Thermo Scientific). 8 seedlings were put in each well. The plant secretions were collected after four days and filtered with a 0.22 µm filter, and stored at 4°C for further use

### Luminescence experiments

Luminescence reporters were grown in either MSgg medium or MSgg medium containing plant secretions. Experiments were carried using a 96 – well plate flat bottom with white opaque walls (Corning). Measurements were performed every 30 min at 30 °C for a period for 24h, using a microplate reader (Synergy 2; BioTek, Winooski, VT, USA). Luciferase activity was normalized to avoid artefacts related to differential cells numbers as RLU/OD.

### Confocal microscopy

Plants cultured with bacteria were washed in PBS and mounted on a microscope slide and covered with a poly-L-Lysine 31 (Sigma)-treated coverslip. Cells were visualized and photographed using a laser scanning confocal microscope (Zeiss LSM 780) equipped with a high resolution microscopy Axiocam camera, as required. Data were captured using Zen black software (Zeiss, Oberkochenm, Germany).

### Flow Cytometer Cell Sorting

Indicated strains used in the experiments were inoculated at an OD of 0.02 in 1.5 ml liquid MSgg without seedlings (control) and MSgg with 14-d-old *A. thaliana* seedlings in a 24-well plate (Thermo Scientific), each well contained four seedling. the set was incubated for 24h in growth chamber (MRC) at 23°C in 12 h light/12 h dark regime. After incubation the seedlings were removed from the liquid medium, washed in PBS and transferred to a 1.5 eppendorf tube in 1 ml of PBS and vortexed for 1 minute, for the purpose of detaching the bacteria from the root. Samples was measured using a BD LSR II flow cytometer (BD Biosciences), using laser excitation of 488 nm, coupled with 505 LP and 525/50 sequential filters.. 100,000 cells were counted and samples were analyzed usingFACS Diva (BD Biosciences).

### Imaging Flow Cytometry

Samples were prepared as for the sorting flow cytometer. Data was acquired by ImageStream X mark II (Luminx corp., Austin Tx) using a 60X lense (NA=0.9). Lasers used were 488nm (200mW) for GFP excitation and 785nm (5mW) for side scatter measurement. During acquisition bacterial cells were gated according to their area (in square microns) and side scatter, which excluded the calilbration beads (that run in the instrument along with the sample). For each sample 100,000 events were collected. Data was analysed using IDEAS 6.3. Single event bacteria were selected according to their area (in square microns) and aspect ratio (width divided by the length of a best-fit ellipse).

### Flow Cytometry

Indicated strains used in the experiments were inoculated at an OD of 0.02 in 1.5 ml liquid MSgg without seedlings (control) and MSgg with 14-d-old *A. thaliana* seedlings in a 24-well plate (Thermo Scientific), each well contained four seedling. The set was incubated for 24h in growth chamber (MRC) at 23°C in 12 h light/12 h dark regime. After incubation the seedlings were removed from the liquid medium, washed in PBS and transferred to a 1.5 Eppendorf tube in 1 ml of PBS and vortexed for 1 minute, for the purpose of detaching the bacteria from the root. Samples was measured using a BD LSR II flow cytometer (BD Biosciences), using laser excitation of 488 nm, coupled with 505 LP and 525/50 sequential filters. For each sample 100,000 cells were counted and samples were analyzed using Diva 8 software (BD Biosciences).

### Imaging Flow Cytometry

Samples were prepared as for the sorting flow cytometer. Data was acquired by ImageStream X mark II (AMNIS, part of Luminex corp., Austin Tx) using a 60X lense (NA=0.9). Lasers used were 488nm (200mW) for GFP excitation and 785nm (5mW) for side scatter measurement. During acquisition bacterial cells were gated according to their area (in square microns) and side scatter, which excluded the calibration beads (that run in the instrument along with the sample). For each sample 100,000 events were collected. Data was analysed using IDEAS 6.3. Single event bacteria were selected according to their area (in square microns) and aspect ratio (width divided by the length of a best-fit ellipse). Focused events were selected by the Gradient RMS and Contrast features (measures the sharpness quality of an image by detecting large changes of pixel values in the image). Cells expressing GFP were selected using the Intensity (the sum of the background subtracted pixel values within the image) and Max Pixel values (the largest value of the background-subtracted pixels) of the GFP channel (Ch02).GFP expression was quantified using the Mean Pixel feature (the mean of the background-subtracted pixels contained in the input mask). The size of bacteria was quantified using the Length feature (measures the longest part of an object, in microns) of the bright-field image.

### Statistical analysis

All experiments were performed three separate and independent times in triplicates, unless stated otherwise. Statistical analyses were performed with GraphPad Prism 9.0 (GraphPad Software, Inc., San Diego, CA).

## Acknowledgments

The Kolodkin–Gal laboratory is supported by the Israel Science Foundation grant number 119/16 and ISF-JSPS 184/20 and Israel Ministry of Science - Tashtiot (Infrastructures) - 123402 in Life Sciences and Biomedical Sciences. IKG is supported by an internal grant from the Estate of Albert Engleman provided by the Angel–Faivovich Fund for Ecological Research, and by a research grant from the Benoziyo Endowment Fund for the Advancement of Science. IKG is a recipient of the Rowland and Sylvia Career Development Chair.

---

**Table S1.**

| Strain | Description | Source or reference |
| --- | --- | --- |
| *B. subtilis* | Wild type | (Branda et al., 2001) |
| *P<sub>pksC</sub>-lux* | *B. subtilis* *sacA::P<sub>pksC</sub>-lux* (Cm<sup>r</sup>), promoter of bacillaene operon tagged to the luciferase reporter integrated in the neutral *SacA* locus | (Ogran et al., 2019) |
| *P<sub>srfAA</sub>-lux* | *B. subtilis* *sacA::P<sub>srfAA</sub>-lux* (Cm<sup>r</sup>), promoter of surfactin operon tagged to the luciferase reporter integrated in the neutral *sacA* locus | (Maan et al., 2021) |
| *P<sub>srfAA</sub>-yfp* | *B. subtilis* *sacA::P<sub>srfAA</sub>-yfp* (Sp<sup>r</sup>), promoter of surfactin operon tagged to the YFP reporter integrated in the neutral *amyE* locus | Avigdor Eldar Lab, TAU, Israel |
| *P<sub>pksC</sub>-gfp* | *B. subtilis* *amyE::P<sub>srfAA</sub>-yfp* (Cm<sup>r</sup>), promoter of bacillaene operon tagged to the GFP reporter integrated in the neutral *amyE* locus | (Ogran et al., 2019) |
| *P<sub>bacA</sub>-gfp* | *B. subtilis* *amyE::P<sub>bacA</sub>-gfp* (Cm<sup>r</sup>), promoter of bacilysin operon tagged to the GFP reporter integrated in the neutral *amyE* locus | (Maan et al., 2021) |
| *P<sub>ppsA</sub>-gfp* | *B. subtilis* *amyE::P<sub>ppsA</sub>-gfp* (Cm<sup>r</sup>), promoter of plipstatin operon tagged to the GFP reporter integrated in the neutral *amyE* locus | (Maan et al., 2021) |
| *P<sub>penP</sub>-gfp* | *B. subtilis* *amyE::P<sub>penP</sub>-gfp* (Sp<sup>r</sup>), promoter of plipstatin operon tagged to the GFP reporter integrated in the neutral *amyE* locus | (Bucher et al., 2019) |

**Cm<sup>r</sup>** – chloramphenicol resistance, **Sp<sup>r</sup>** – spectinomycin resistance

